# Measuring the gut microbiome in birds: comparison of faecal and cloacal sampling

**DOI:** 10.1101/160564

**Authors:** Elin Videvall, Maria Strandh, Anel Engelbrecht, Schalk Cloete, Charlie K. Cornwallis

**Author notes:** Corresponding author; Elin Videvall.

## Abstract

The gut microbiomes of birds and other animals are increasingly being studied in ecological and evolutionary contexts. While methods for preserving samples and processing high-throughput sequence data to characterise bacterial communities have received considerable attention, there has been little evaluation of non-invasive sampling methods. Numerous studies on birds and reptiles have made inferences about gut microbiota using cloacal sampling, however, it is not known whether the bacterial community of the cloaca provides an accurate representation of the avian gut microbiome. We examined the accuracy with which cloacal swabs and faecal samples measure the microbiota in three different parts of the gastrointestinal tract (ileum, caecum, and colon) using a case study on juvenile ostriches, *Struthio camelus*, and high-throughput 16S rRNA sequencing. We found that faeces were significantly better than cloacal swabs in representing the bacterial community of the colon. Cloacal samples had a higher abundance of Gammaproteobacteria and fewer Clostridia relative to the gut and faecal samples. However, both faecal and cloacal samples were poor representatives of the microbial communities in the caecum and ileum. Furthermore, the accuracy of the sampling methods in measuring the abundance of different bacterial taxa was highly variable: Bacteroidetes was the most highly correlated phylum between all three gut sections and both methods, whereas colonic Actinobacteria correlated strongly only with faecal samples. This study demonstrates that sampling methods can have significant effects on the inferred gut microbiome in studies of birds. Based on our results, we recommend sampling faeces, whenever possible, as this provides the most accurate assessment of the colon microbiome. The fact that neither sampling technique portrayed the bacterial community of the ileum or the caecum illustrates the difficulty in non-invasively monitoring gut bacteria located further up in the gastrointestinal tract. These results have important implications for the interpretation of avian gut microbiome studies.

## Introduction

The community of bacteria harboured within the gastrointestinal tract of animals – ‘the gut microbiome’ – has been established as an important determinant of host health and physiology (Sekirov *et al.* 2010). Although research has largely focused on humans and model organisms, it is becoming increasingly recognised that the gut microbiome may play an important role in a variety of ecological and evolutionary processes, as it has been associated with disease resistance, behaviour, mate selection, longevity, and adaptation (Sharon *et al.* 2010; Koch & Schmid-Hempel 2011; Muegge *et al.* 2011; Ezenwa *et al.* 2012; Brooks *et al.* 2016; Smith *et al.* 2017). As a result, it is necessary that accurate methods for monitoring the gut microbiome in ecologically relevant contexts are developed. To date, multiple studies have focused on the reliability of methods for storing and preserving samples, as well as techniques for processing data from high-throughput sequencing (see e.g., Debelius *et al.* 2016; Song *et al.* 2016). However, it remains unclear whether different sampling techniques accurately represent the bacterial communities in different parts of the gastrointestinal tract.

A large number of studies investigating the gut microbiome of birds and reptiles have sampled bacteria from the cloaca (Bowman & Jacobson 1980; Cooper *et al.* 1985; Lombardo *et al.* 1996; D’Aloia *et al.* 1996; Mills *et al.* 1999; Dickinson *et al.* 2001; Lamberski *et al.* 2003; Moreno *et al.* 2003; Lucas & Heeb 2005; Maul *et al.* 2005; Santoro *et al.* 2006; Hoar *et al.* 2007; Klomp *et al.* 2008; Ruiz-Rodríguez *et al.* 2009a; b; Martin *et al.* 2010; Xenoulis *et al.* 2010; Santos *et al.* 2012; Charruau *et al.* 2012; Dewar *et al.* 2013, 2014; van Dongen *et al.* 2013; Stenkat *et al.* 2014; Allegretti *et al.* 2014; Matson *et al.* 2015; Stanley *et al.* 2015; Kreisinger *et al.* 2015; Barbosa *et al.* 2016; Merkeviciene *et al.* 2017; Lobato *et al.* 2017; Ganz *et al.* 2017). Cloacal sampling is widely used because it is straightforward to perform, allows repeated sampling of individuals, and the possibility of reliably obtaining samples from all individuals at the same time. This can provide practical advantages over faecal sampling, which may be unreliable and provides potential difficulties in identifying sample ownership and time of defecation.

It is, however, not known if the microbiota of the cloaca provides an accurate reflection of the bacterial community in the gut, and whether cloacal sampling is a good alternative method to faecal sampling. From a theoretical point of view, there are reasons to believe that the bacterial community of the cloaca is not simply seeded with bacteria from faeces. The cloaca constitutes the single posterior opening for the digestive, reproductive, and urinary tract in birds, reptiles, amphibians, sharks, rays, and a few mammals, and as such represents an important barrier to foreign bodies, including pathogens. For example, during copulation, many bird species engage in a so-called “cloacal kiss”, where they exchange not only sperm, but also cloacal microbes (Kulkarni & Heeb 2007; White *et al.* 2010). In fact, the avian cloaca has a specialised immune organ, the bursa of Fabricius, that is involved in the development of B lymphocytes and antibody production (Warner & Szenberg 1964), and enables contact between cloacal microbes and the lymphoid system (Schaffner *et al.* 1974). Furthermore, the cloacal mucosa likely constitutes an environment that is mostly aerobic compared to the anaerobic environment of the gastrointestinal lumen, as this is the case for the mammalian rectum (Albenberg *et al.* 2014; De Weirdt & Van de Wiele 2015). Taken together, the proximity of the mucosal cloacal microbiome to both the external environment and host tissue, including secreted mucus with immune cells and antimicrobial molecules, likely results in a microbial environment different from that of the gut, and potentially therefore structural differences in microbiota. Nevertheless, several studies investigating bacterial gut composition in birds directly refer to cloacal swabs as faecal samples, with the assumption that they are equivalent (Dewar *et al.* 2013, 2014; Allegretti *et al.* 2014; Stanley *et al.* 2015).

In line with the idea that the cloaca may accommodate different bacteria, two studies evaluating cloacal swabs with caecal samples in chickens found large differences in bacterial communities (Stanley *et al.* 2015; Zhang *et al.* 2017). It has been argued, however, that cloacal samples may still reflect the presence of the vast majority of caecal bacteria if they are sequenced deep enough (Stanley *et al.* 2015), and it is unclear whether faecal sampling would provide a more accurate picture. This raises the issue of whether particular sampling techniques are superior at measuring specific groups of bacteria in the gut microbiome. For example, certain bacterial taxa may be more widely distributed along the gastrointestinal tract and hence easier to monitor, while other taxa may be confined to specific locations in the gut and thus not well represented by any sampling method. Uncovering what attributes of the gut microbiome different types of sampling methods are able to measure, and what they can infer about the microbial communities present in the different sections of the intestinal tract will be essential to advance our understanding of host microbiomes.

In this study we evaluate the accuracy of two commonly used microbiome sampling techniques for birds: cloacal swabs and faecal samples. We test the similarity of the cloacal and faecal microbiomes to three parts of the gastrointestinal tract: ileum, caecum, and colon. For this purpose, 20 juvenile ostriches between four to six weeks old were used as a case study.

## Materials and methods

### Study species

We used the ostrich, *Struthio camelus*, as case study species, kept under controlled conditions at the Western Cape Department of Agriculture ostrich research facility in Oudtshoorn, South Africa. The samples in this study were obtained in 2014 from a total of 20 juveniles, which included ten individuals four weeks old and ten individuals six weeks old. Ostrich chicks can easily be maintained and handled in an experimental setting, and this specific age group is ideal in size and temperament for both faecal sampling and dissection, allowing us to efficiently retrieve all necessary samples in a standardised way. The chicks were housed and reared with their contemporaries in four separate groups in indoor pens in the same building, containing approximately 35-40 individuals in each group at the time of sampling. During the daytime they had access to outside enclosures where they could peck freely in soil, and were given ad libitum access to fresh water and food throughout the trial.

### Sample collection

Faecal samples were collected from all chicks one day before scheduled euthanization and dissection, by placing sterile plasters over their cloaca and retrieving the collected fresh faeces approximately one hour later. Two to three chicks were randomly selected from each group for gut sampling, totalling ten individuals per sampling event, one at four weeks of age and one at six weeks. Before dissection, the 20 randomly selected chicks were euthanized by a licensed veterinarian who severed the carotid artery. All procedures were approved by the Departmental Ethics Committee for Research on Animals (DECRA) of the Western Cape Department of Agriculture, reference number R13 / 90. During the dissection we collected four samples from each individual: cloacal swabs and samples from the ileum, caecum, and colon. Cloacal samples were collected by using sterile cotton swabs that were briefly moistened in phosphate-buffered saline (PBS), and the tip carefully inserted in the cloaca of the birds and gently rotated.

To minimize contamination between samples and individuals, a number of precautions were taken. Lab benches and surfaces were routinely sterilized with 70% ethanol, and equipment used during the dissection was first cleaned with hot water, then rinsed with 70% ethanol and subsequently placed in the open flame of a Bunsen burner between each sample collection for sterilization. Control swabs were collected during both dissection events and during the faecal sampling. The control swabs followed the same initial procedure as the cloacal swabs (dipping sterile cotton swabs in PBS), but instead of sampling the bird, they were exposed to potential microbes in the air by waving the wet swab around in the dissection / sampling room. All samples were collected in plastic 2 ml micro tubes (Sarstedt, cat no. 72.693) between October 28 and November 12, 2014, and stored at-20 °C within two hours of collection. They were subsequently transported on ice to a laboratory and stored at-20 °C.

### DNA isolation, library preparation, and amplicon sequencing

We prepared sample slurries for all sample types with guidance from Flores *et al.* (2012) and subsequently extracted DNA using the PowerSoil-htp 96 well soil DNA isolation kit (MoBio Laboratories, cat no. 12955-4) as recommended by the Earth Microbiome Project (http://www.earthmicrobiome.org) (for full details please see Supplementary Methods available online). Libraries for sequencing of the 16S rRNA V3 and V4 regions were prepared using the primers Bakt_341F and Bakt_805R (Herlemann *et al.* 2011) according to the Illumina 16S Metagenomic Sequencing Library Preparation Guide (Part # 15044223 Rev.B). All samples in this study (Table S1) were sequenced in one 300-bp paired end run on an Illumina MiSeq platform at the DNA Sequencing Facility, Department of Biology, Lund University, Sweden. In a subsequent run, we sequenced blank samples and additional control samples that were collected during the trial for a related project. These control samples were not essential for this particular study, but were included to increase the number of controls. As a result, a total of 117 different samples plus 54 sample replicates (see Supplementary Methods) were part of this study.

### Data processing

The 16S amplicon sequences were quality controlled using FastQC (v. 0.11.5) (Andrews 2010) together with MultiQC (Ewels *et al.* 2016). Primers were removed from the sequences using Trimmomatic (v. 0.35) (Bolger *et al.* 2014) and the forward reads were retained for analyses. Quality filtering of the reads were executed using the script multiple_split_libraries_fastq.py from QIIME (v. 1.9.1) (Caporaso *et al.* 2010). All bases with a Phred score < 25 at the 3’ end of reads were trimmed and samples were multiplexed into a single high-quality multi-fasta file.

Operational taxonomic units (OTUs) were assigned and clustered using Deblur (v. 1.0.0) (Amir *et al.* 2017). Deblur circumvents the problems surrounding clustering of OTUs at an arbitrarily threshold by obtaining single-nucleotide resolution OTUs after correcting for Illumina sequencing errors. This results in exact sequence variants (ESVs), also called amplicon sequence variants (ASVs), oligotypes, and sub-OTU (sOTUs). In order to avoid confusion, we chose to call these units OTUs, but the reader should be aware that they differ from the traditional 97% clustering approach (Callahan *et al.* 2017). The minimum reads-option was set to 0 to disable filtering inside Deblur, and all sequences were trimmed to 220 bp. We used the biom table produced after both positive and negative filtering, which by default removes any reads which contain PhiX or adapter sequences, and only retains sequences matching known 16S sequences. Additionally, PCR-originating chimeras were filtered from reads inside Deblur (Amir *et al.* 2017).

Taxonomic assignment of OTUs was performed using the Greengenes database (DeSantis *et al.* 2006). We filtered all samples on a minimum read count of 1000 sequences, resulting in three out of 171 samples being excluded (one ileum and two control samples). We further filtered all OTUs that only appeared in one sample, resulting in the removal of 8,846 OTUs, with 3,015 remaining. The samples with technical replicates (two control samples and seven individuals with replicates for all sample types; see Supplementary Methods) had the replicates merged within their respective sample type (i.e. ileum.rep1 + ileum.rep2) to increase the amount of valuable sequence information. The analyses were evaluated with both rarefied and non-rarefied data, which produced extremely similar and comparable results. We therefore present the results from the non-rarefied data in this study, as recommended by McMurdie & Holmes (2014).

### Data analyses

All analyses were performed in R (v. 3.3.2) (R Core Team 2017). We calculated alpha diversity using the Shannon measure with absolute abundance of reads, and distance measures with the Bray-Curtis distance method on relative read abundances in phyloseq (v. 1.19.1) (McMurdie & Holmes 2013). Differences between the microbiota of cloacal and faecal samples to the microbiota of each gut section were examined using permutational multivariate analysis of variances (PERMANOVA) on Bray-Curtis distances using the Adonis function in vegan (v. 2.4-2) with 1000 permutations (Oksanen *et al.* 2017). To analyse if there were differences in the variance (dispersion) between sample types, we used the multivariate homogeneity of group dispersions test (betadisper) in vegan (Oksanen *et al.* 2017), followed by the Tukey’s ‘Honest Significant Difference’ method. Blank and control samples showed highly dissimilar microbial composition to all other sample types (see Figures S1, S2, S3, and S4) and were not included in any further analyses.

To evaluate bacterial abundances, we first filtered out all OTUs with less than 10 sequence reads and then, using DESeq2 (v. 1.14.1), counts were modelled with a local dispersion model and normalised per sample using the geometric mean (see the DESeq2 manual) (Love *et al.* 2014). We examined the strength of the correlations between the abundance of bacteria (normalised OTU abundance), both at the level of phylum and class, in the three parts of the gut in relation to the abundances in both cloacal swabs and faecal samples. Two sets of correlations were performed, one where each data point represented the mean number of OTUs in that bacterial taxon averaged across the 20 individuals (‘correlations across bacteria’, n = number of OTUs per bacteria phylum or class) and one set of correlations where each data point represented the abundance of a bacterial taxon per individual (‘correlations across individuals’, n = 20). We used Spearman’s rank-order correlation and tested the differences between correlations obtained for cloacal samples and those from faecal samples using cocor (v. 1.1-3) (Diedenhofen & Musch 2015).

Differential abundances between sample types were subsequently tested in DESeq2 with a negative binomial Wald test using individual ID as factor and with the beta prior set to false (Love *et al.* 2014). The results for specific comparisons were extracted (e.g. faeces versus ileum) and p-values were corrected with the Benjamini and Hochberg false discovery rate for multiple testing. OTUs were labelled significant if they had a corrected p-value (q-value) < 0.01. Plots were made using phyloseq (McMurdie & Holmes 2013) and ggplot2 (Wickham 2009).

## Results

### Overall microbiome composition in different sample types

First, we evaluated the overall pattern of the microbial community reflected by the two sampling techniques (cloacal swabs and faeces) and the three different sections of the avian gastrointestinal tract (Figure 1). The abundance of bacterial taxa in the microbiomes of the caecum, colon, and faeces showed large overall similarities (Figures 1C, 1D), especially the faecal and colon samples which closely clustered in the network plot (Figure 1B). These three sample types also had the highest and most similar alpha diversity values (colon mean Shannon’s diversity index H = 4.47, faeces H = 4.25, and caecum H = 4.16; Figure 1A). Bacteria from the classes Clostridia (phylum: Firmicutes) and Bacteroidia (phylum: Bacteroidetes) mainly dominated in the caecum, colon, and faeces (Figures 1C, 1D). In contrast, the cloacal and ileum samples showed large overall taxa dissimilarities in microbiota composition compared to the other samples types (Figure 1). The microbiome of the cloaca had significantly lower alpha diversity compared to the caecum, colon, and faeces (H = 3.40, paired Wilcoxon signed rank test against caecum: V = 37, p = 0.009; against colon: V = 4, p < 0.0001, and against faeces: V = 17, p = 0.0004), and so did the ileum (H = 2.50, pairwise comparisons against caecum, colon, and faeces: V = 0, p < 0.0001). The cloaca showed a distinct microbial community from all other samples at the class level with a high relative abundance of Gammaproteobacteria and Bacilli, and a lower abundance of Clostridia (Figures 1C, 1D). The ileum also showed higher abundance of Bacilli and lower abundance of Clostridia, but was overall dissimilar to all other samples with a high representation of Betaproteobacteria and very few Bacteroidia (Figures 1C, 1D).

**Figure 1.**
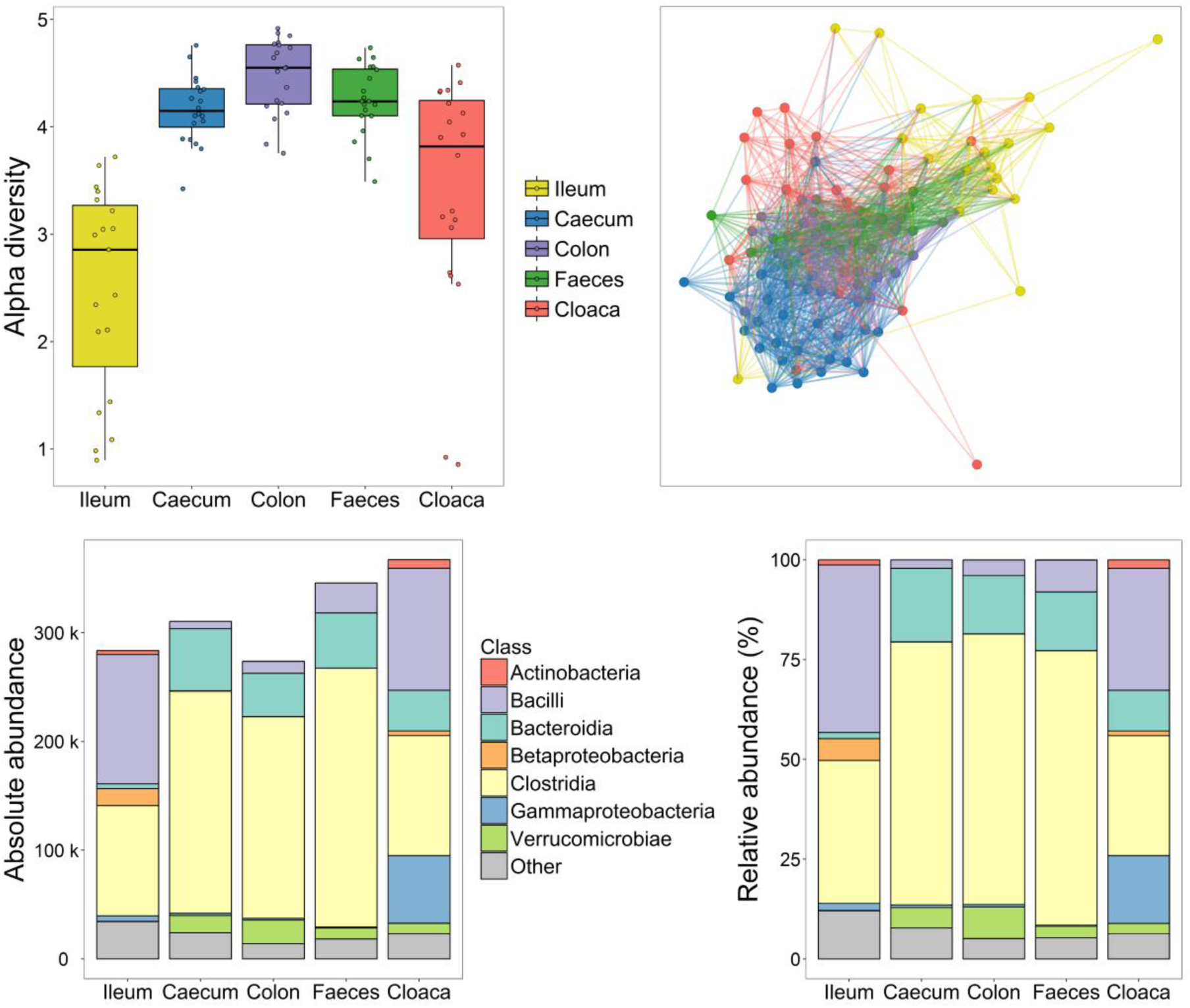
Overall microbiota similarities and differences between sample types. (A) Boxplot of Shannon alpha diversity within sample types. (B) Network of Bray-Curtis distances between samples where colours indicate sample type and lines are drawn to nearest neighbours with a maximum distance of 0.85. The barplots show the (C) absolute and (D) relative abundance of all OTUs for each sample type. Taxonomic classifications are coloured at the class level.

### Distances between the microbiomes of the cloaca and faeces to the gut sections

Second, to evaluate overall microbiota distance dissimilarities between the two sample methods to the gut samples, we conducted multivariate analyses of variance (Adonis). All comparisons were significantly different from each other (PERMANOVA: p < 0.001), indicating that each sample type harbours a unique microbiome. This was due to differences in mean distances between communities, not differences in variances, as there was no difference in dispersion between sample types (multivariate homogeneity test of group dispersions: adjusted p > 0.152). The two most similar sample types were the faeces and colon, which resulted in a low R^2^ (0.069), whereas the cloaca and colon were more dissimilar (R^2^ = 0.099). Both sampling methods reflected greater dissimilarities to the gut sections further up in the gastrointestinal tract, with faecal samples being more distant to the caecum (R^2^ = 0.191) and the ileum (R^2^ = 0.160), as were cloacal samples (caecum: R^2^ = 0.136, ileum: R^2^ = 0.145).

To directly test how well cloacal swabs and faecal samples represented the microbiota in the gut, we calculated community Bray-Curtis distances between the faecal and cloacal samples to each of the three sections of the gut for each individual (Figure 2). Neither sampling technique was particularly good at measuring the microbiome of the ileum (cloacal mean distance = 0.87, faecal mean distance = 0.84, paired Wilcoxon signed rank test: V = 136, p = 0.104) or the caecum (cloacal mean distance = 0.82, faecal mean distance = 0.84, V = 88, p = 0.546). However, the faecal samples were significantly closer in distance to the colon than the cloacal samples were (cloacal mean distance = 0.74, faecal mean distance = 0.63, V = 164, p = 0.027) (Figure 2).

**Figure 2.**
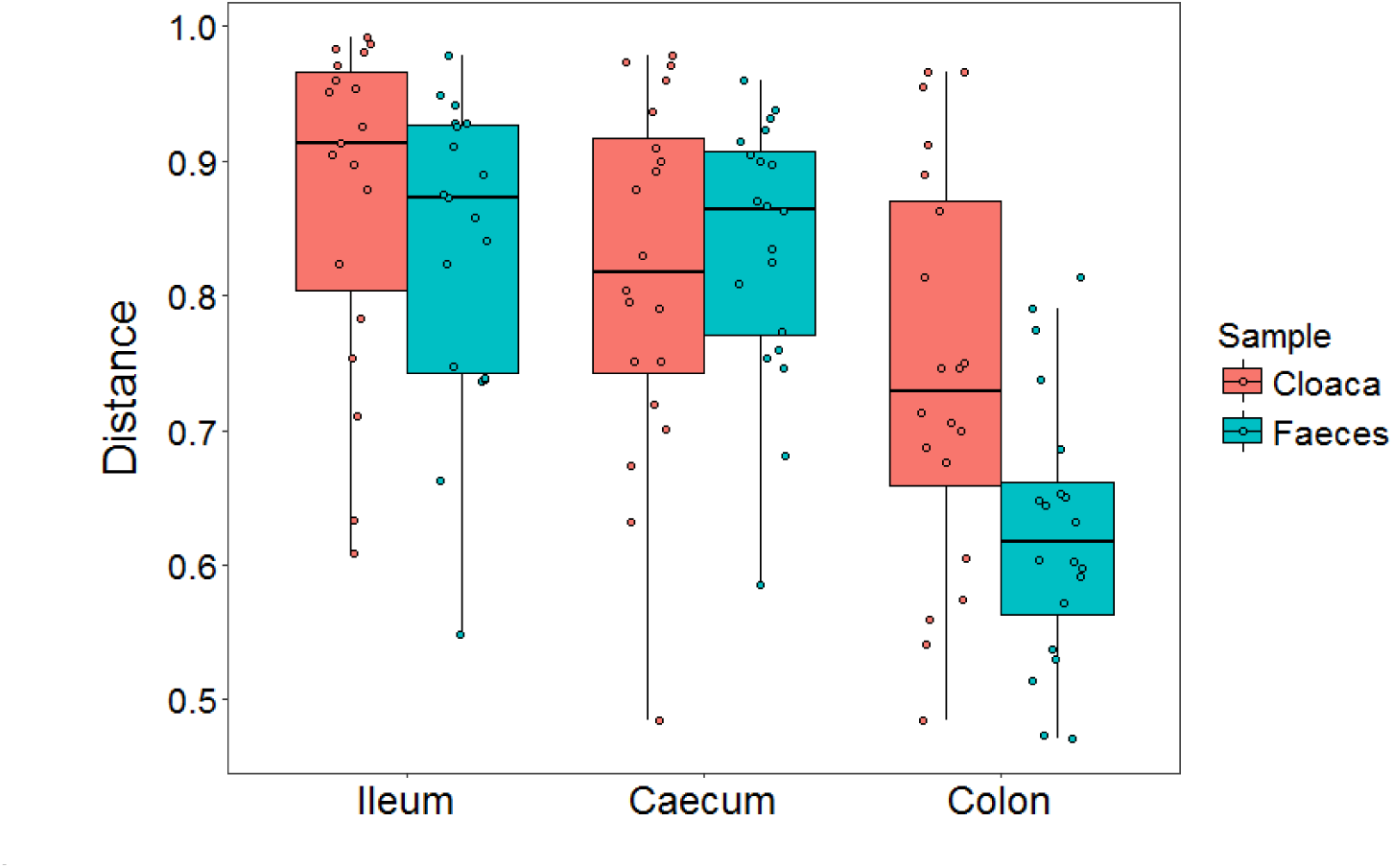
Bray-Curtis distances between the microbiota in cloacal and faecal samples compared to the microbiota in three parts of the gut (ileum, caecum, colon) within the same host individual. Higher distance measures indicate higher dissimilarity, where 1 = completely dissimilar bacterial community.

### Correlation of bacterial abundances in the cloaca and faeces with the gut sections

We further evaluated how accurately the sampling techniques represented the abundance of all OTUs in the ileum, caecum, and colon, and found that the correlations of both faecal and cloacal samples with the ileum and caecum were weak (r = 0.045 – 0.268; Figure 3). Conversely, the correlations with the colon were strong, especially for the faecal samples (r = 0.558 versus r = 0.476 for cloacal swabs; Figure 3). When analysing the abundances of different bacterial phyla separately, we again found that the correlations between the sampling methods and the ileum were weak for all six phyla (r < 0.275; Table S2). The phyla abundance correlations were stronger for the colon (r = 0.246 – 0.803; Table S2), but highly variable for the caecum (r =-0.127 – 0.633; Table S2). Similar patterns of correlation were also found when analysing abundances across different bacterial classes (Table S3). More specifically, the phylum Bacteriodetes had the strongest correlations between both sampling methods and each of the three gut sections (Table S2), and at a lower taxonomic level, the two classes Bacteriodia (phylum: Bacteroidetes) and Coriobacteriia (phylum: Actinobacteria) displayed strong correlations between each of the two sampling techniques to both the caecum and colon (Table S3). Overall, the correlations between faecal samples and cloacal swabs to the different parts of the gut were similar with a few exceptions. For example, the abundance of Actinobacteria in the colon and caecum appeared to be better represented in faeces, whereas the abundance of Tenericutes and Betaproteobacteria in the same intestinal regions appeared to be better represented in cloacal swabs.

**Figure 3.**
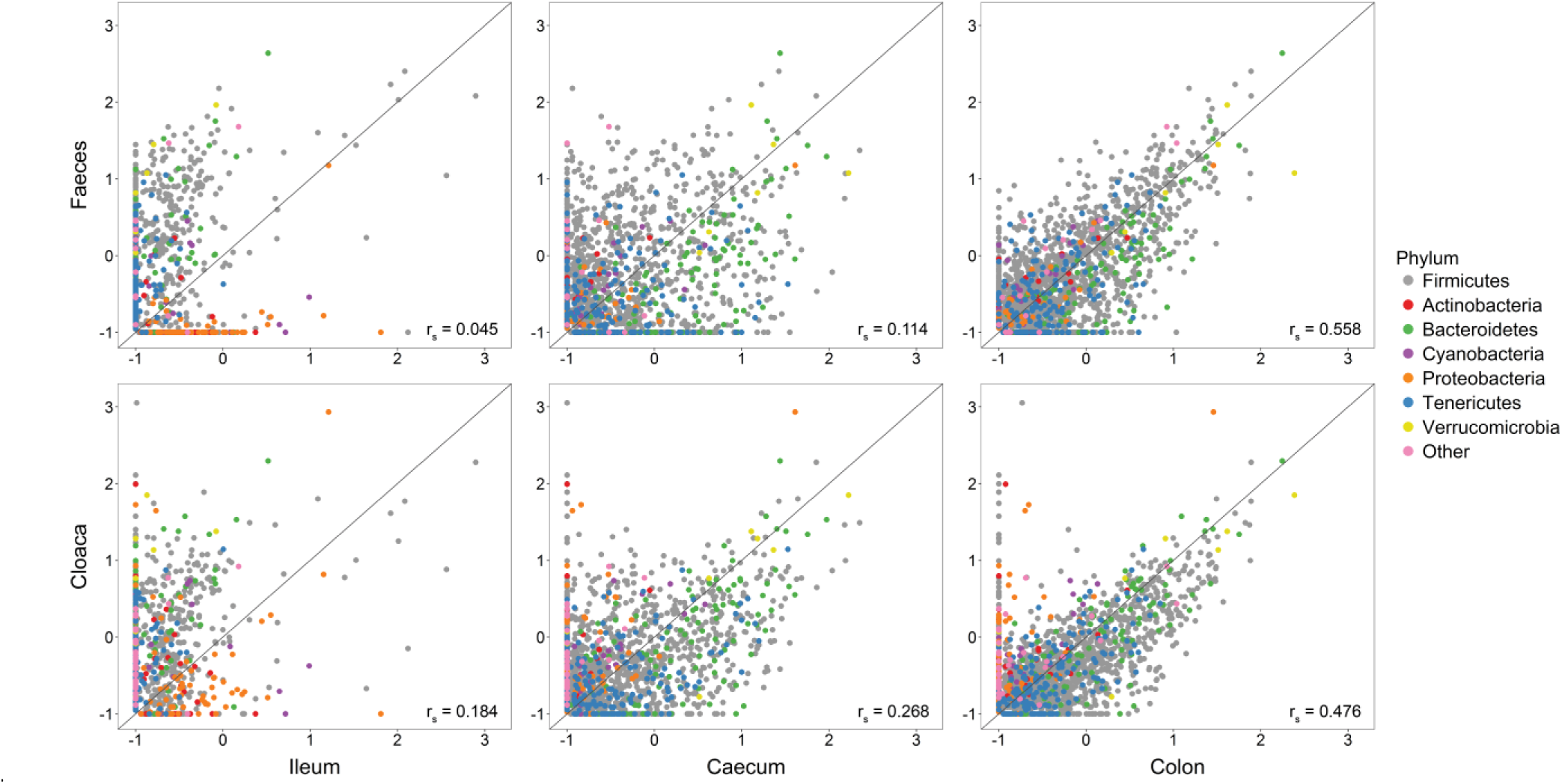
Correlations in OTU abundances between each of two sampling methods (cloaca and faeces) compared to three sections of the gut (ileum, caecum, and colon). OTU abundances were normalised according to the DESeq2 method and log-transformed (+ 0.1) for graphical purposes. The straight line shows the 1:1 relationship between the samples, and the r_s_ values display the Spearman rank correlation across all OTUs.

In addition, we examined how well the abundances of different bacteria correlated between samples from the same host individuals (Figure S4). Overall OTU abundance in the ileum was weakly correlated with faeces (r = 0.162), but more strongly with cloacal swabs (r = 0.493). In contrast, individual faecal samples showed extremely high correlations to both the caecum (r = 0.872) and the colon (r = 0.893), whereas cloacal samples only showed intermediate correlations to the caecum (r = 0.509) and colon (r = 0.538) (Figure S4).

### Differences in abundance of specific OTUs in the cloaca and faeces versus the gut sections

Next, we analysed whether specific OTUs were more or less abundant when using either of the two sampling techniques by testing for significant differences (q < 0.01) in OTU abundance in the cloacal and faecal samples compared to the three gut sections (Figure 4; Tables S4-S9). Consistent with our previous analyses, we found the highest number of significantly different OTUs when comparing the ileum to both the faecal (n = 307) and cloacal samples (n = 250), followed by the comparisons with the caecum (144 significant OTUs for faeces versus 123 for cloacal swabs). The colon showed the least differences in abundance to both sampling methods, but the cloacal samples had twice as many significant OTUs (n = 64) compared to faecal samples (n = 32), indicating substantially more differences between cloaca-colon than faeces-colon (Tables S8-S9).

**Figure 4.**
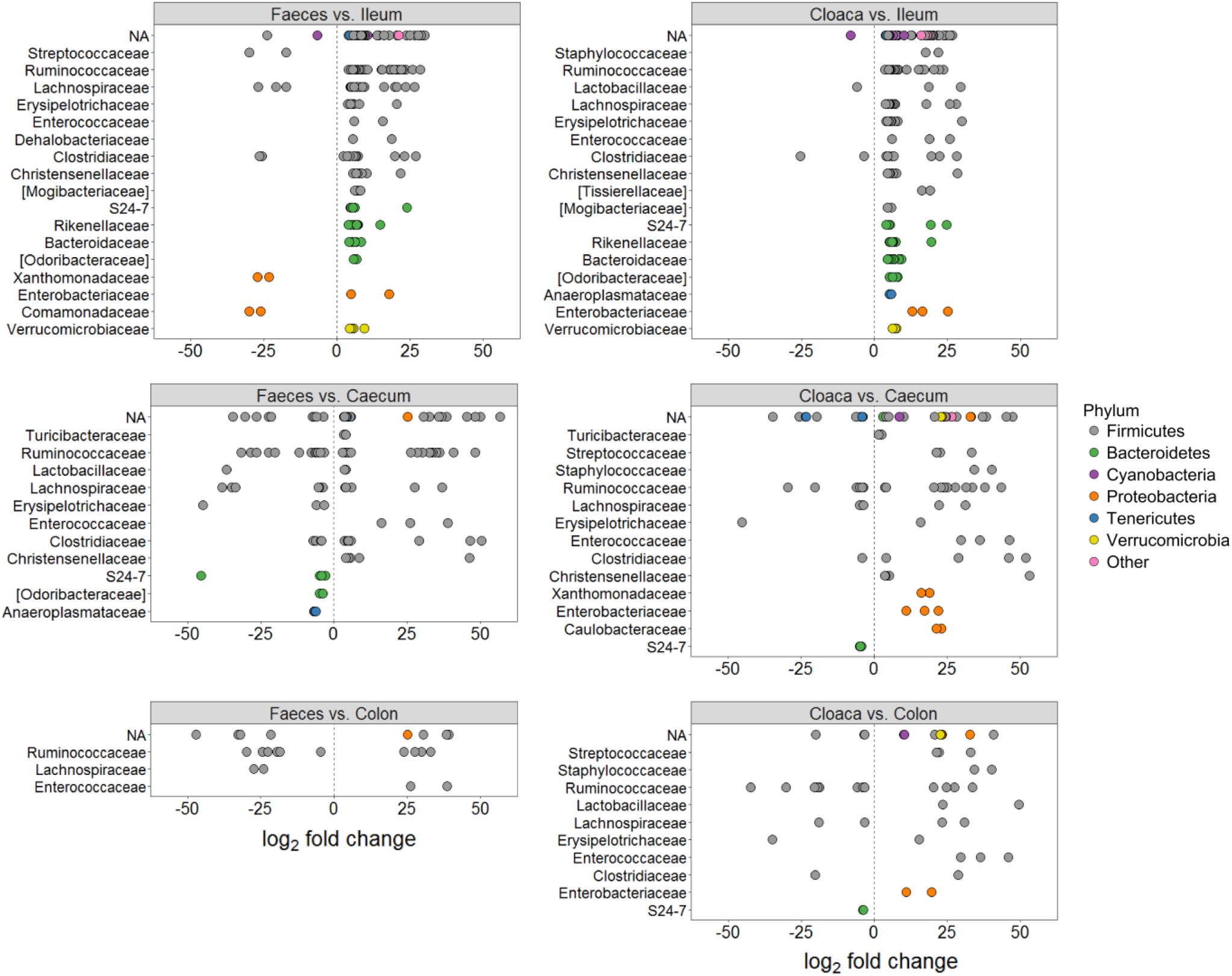
OTUs that were significantly different in abundance in the sampling methods (faeces and cloaca) relative to three parts of the gut (ileum, caecum, and faeces). The y-axes show taxonomic families and all OTUs have been coloured within their respective phylum. Positive log2 fold changes signify increased OTU abundance in either faeces (left column) or cloacal swabs (right column), and negative log2 fold changes display increased abundance in one of the gut sections (ileum, caecum, or colon). Families with only a single significant OTU are not shown due to space limitations; complete dataset can be found at Tables S4-S9. Family names in brackets are proposed taxonomies by Greengenes.

We further evaluated the taxa that showed significantly different abundances across the six sample comparisons. Relative to the ileum, a large number of OTUs in the phylum Firmicutes were significantly more abundant in both faeces and the cloaca (Figure 4; Tables S4-S5). The most significant Firmicutes families included Ruminococcaceae, Lachnospiraceae, Erysipelotrichaceae, Clostridiaceae, and Christensenellaceae (Tables S4-S5). The Enterobacteriaceae (Proteobacteria), the Verrucomicrobiaceae (Verrucomicrobia), and several families within the Bacteroidetes were also significantly more abundant in both faeces and the cloaca compared to the ileum. When comparing sampling methods against the caecum, several Firmicutes bacteria showed significantly different abundances in both directions (Figure 4). The caecum showed, however, a significantly higher abundance of Bacteroidetes relative to both the cloaca and faeces, with one exception: an OTU within the Rikenellaceae, which was completely absent in the caecum samples but present in both sampling methods. Interestingly, the cloaca had a lot more significantly different Proteobacteria OTUs (n = 19) than faeces did (n = 2) in the comparison with the caecum, and 94.7% of those were more abundant in the cloacal samples (Figure 4; Tables S6-S7). Finally, the comparison between the colon and faeces only resulted in 13 significantly different bacterial families within five phyla, while the difference between the colon and cloaca was much larger and phylogenetically broader, representing 28 significantly different families from 11 phyla (Figure 4; Tables S8-S9).

## Discussion

Measuring the gut microbiome of birds and other animals is becoming increasingly important for ecologists and evolutionary biologists due to its potential implications for host fitness. Numerous studies sample either the cloacae or faeces of birds as a proxy for estimating the bacterial community in the gut, however, it has remained untested whether cloacal or faecal sampling constitute accurate ways of measuring avian gut bacteria. In this study we examined the microbiota of cloacal swabs and faeces and compared them to the microbiota in three different sections of the gastrointestinal tract. We found that cloacal swabs were less accurate at representing the microbiome of the colon relative to faecal samples, which had more similar community composition and abundances of bacteria. Neither faeces nor cloacal swabs could, however, accurately estimate the bacterial communities of the ileum and the caecum. These results have important implications for the interpretation of bird gut microbiomes, and we hope they will aid researchers in the planning of future studies.

The different sections of the gastrointestinal tract were associated with spatial heterogeneity in their bacterial composition, which is largely expected given their different physiological functions. The ileum is the final part of the small intestine and has a primary role of absorbing nutrients from food while maintaining a neutral pH. In our study, the ileal microbiome had the lowest alpha diversity, which is consistent with other studies investigating the small intestine of birds and reptiles (Bjerrum *et al.* 2006; Danzeisen *et al.* 2015; Kohl *et al.* 2017). It also had the highest relative abundance of Bacilli and Betaproteobacteria compared to the other sample types. The second sample site of the gastrointestinal tract, the caecum, provides important functions by breaking down plant and fibrous material, and birds typically have two caeca, located between the small and large intestines. Although the caecal samples in our study were dissimilar to other sample types, they most closely clustered with samples from the colon, at least at higher taxonomic levels. Both of these intestinal regions had high abundances of Clostridia and Bacteroidia. In comparison to both faecal and cloacal samples, the caecum had a significantly higher abundance of several Bacteroidetes, similar to previous research on the chicken caecum (Stanley *et al.* 2015). The final part of the intestinal tract, the colon, has a primary function to absorb water and salt from ingested material. The colon samples in our study had the highest alpha diversity of all sample types and the strongest taxa correlations to both sampling methods, although significantly better to faeces than to the cloaca.

The similarities of both the cloacal and faecal microbiota to that of the gut increased the further down the gastrointestinal tract we sampled, as perhaps expected given the proximity to the cloaca. Nevertheless, there was substantial variation in how well bacterial abundances in different parts of the gut correlated with faecal and cloacal samples across different taxonomic groups, both when examining across individual OTUs and across individual hosts. This variation does not appear to be simply explained by differences in the total abundance of different bacteria (e.g. more abundant bacteria might be more widely distributed in the gut and so more strongly correlated across samples), as some classes of bacteria had high numbers of OTUs, but were poorly correlated and vice versa (Table S3). The strength of correlations between different sample types may potentially reflect the fact that different bacteria vary markedly in the environmental conditions they can tolerate, and hence the breadth of their spatial distribution in the gut. The causes underlying this variation require further investigation, but by presenting effect sizes of the strength of associations we hope to provide useful information on which bacteria can reliably be monitored in different locations of the gastrointestinal tract (Tables S2-S3).

A common goal of microbiome studies, particularly in ecological contexts, is to understand how gut bacteria relate to phenotypic variation. Because it is not feasible to collect intestinal samples in wild birds without highly invasive techniques, faecal or cloacal samples are often the only option, especially if repeated sampling is required. Our results suggest that the bacterial communities in the upper and middle gastrointestinal tract are distinct from those recovered by non-invasive sampling methods, and as such, any inferences made about the gut microbiome and its relationship to phenotypic variation may only be possible for processes occurring in the colon. Further studies are needed to investigate if the results of this study hold true for other animals with cloacae, such as frogs, lizards, and egg-laying mammals. Most mammals possess a rectum instead of a cloaca, which differs in both function and physiology, and rectal swabs are therefore likely to differ substantially to cloacal swabs in the degree to which they are useful for monitoring gut microbiomes. The current evidence as to whether rectal swabs constitute a representative sampling method of the gut microbiome of mammals is conflicting and suffers from low sample sizes, thus warranting additional evaluation (Budding *et al.* 2014; Alfano *et al.* 2015; Bassis *et al.* 2017).

In conclusion, for gut microbiome sampling of birds, we recommend faecal samples whenever possible, as this sampling procedure best captured the bacterial community of the colon.

## Acknowledgements

We are grateful to all the staff at the Oudtshoorn Research Farm, Western Cape Government and Naomi Serfontein, Maud Bonato, and Julian Melgar for assisting during the collection of samples. Adriaan Olivier, Klein Karoo International, generously provided instructions on dissection and performed euthanization of chicks. Thomas Johansson performed the sequencing and Paul McMurdie, Michiel Op De Beeck, Se Jin Song, and Amnon Amir provided valuable advice on analyses. This research was partially funded by research grants to E.V. from the Helge Ax:son Johnson Foundation, the Längmanska Cultural Foundation, the Lund Animal Protection Foundation, the Lars Hierta Memorial Foundation, and the Royal Physiographic Society of Lund. It was further funded by a Wallenberg Academy Fellowship and VR grant to C.K.C. and by the Western Cape Government.

## Author contributions

Study design was planned by E.V. and C.K.C. Sample collection was performed by E.V., C.K.C., and A.E. Animal facilities were provided by S.C., and supervised by A.E. The laboratory work was planned and performed by M.S. The analyses were performed by E.V. with C.K.C. assisting. E.V. wrote the paper with input from all authors.

## Supporting information

Supporting information has been made available online.

### Appendix S1

Supplementary Methods, Supplementary Figures S1 – S4, Supplementary Tables S1 – S3, and Supplementary References.

### Appendix S2

Supplementary Tables S4 – S9.

